# Forecasting novel therapeutic development in biomedical research

**DOI:** 10.64898/2026.05.29.728775

**Authors:** Salsabil Arabi, B. Ian Hutchins

## Abstract

**Background:** Early identification of promising drug research topics is challenging yet crucial for the scientific community to accelerate the development of novel therapeutics. In this work, we leverage large-scale data from the biomedical literature to extract predictive features to identify promising therapeutic research topics at an early stage.

**Methods:** We use a modularity maximizing machine learning algorithm to divide the global citation graph of biomedical literature into a time series of research topic clusters. We extract features for predictive analytics based on citation activity, publication content, and measurable flocking of scientists into novel research topics.

**Results:** Here we report a maximum likelihood model that prospectively identifies research topics culminating in the studies that were pivotal for 99 Food and Drug Administration (FDA)-approvals for drugs (F1-score of 0.84). 80% of target drugs are predicted in advance, with 65% predicted 8 or more years before approval. This predates the start of phase 2 clinical trials in the vast majority of positive predictions.

**Conclusions:** These results show this approach can efficiently flag research topics generating approved drugs several years prior to approval using public data that would have been contemporaneous at the time of prediction. Thus, reliable forecasting can be accomplished with a high-level view of the publication and citation behavior of scientists, without depending on clinical trial data that may only be deposited with a significant lag. This demonstrates that it is possible to detect early signals of future FDA approved therapies even without any specialized information about these applied research efforts.

**Plain Language Summary:** Many people (such as patients, doctors, and scientists) want to know when a new disease treatment is likely to become available. We developed a method that tracks research topics as they emerge and mature to predict when a new disease treatment is likely. Our computational model predicts, 8 or more years ahead of approval, which topics are likely to give rise to a new disease treatment. It does not predict which drug will be approved, but does predict which disease will have a new treatment. In addition to giving patients and doctors an indication of treatment changes, this can be used by scientific organizations and funders to identify promising research topics and target financing into them in an effort to accelerate disease treatment.

**Teaser:** Large-scale data analysis can use the full set of scientific citations to predict which areas of research will yield new FDA approved drugs, years in advance.

## Introduction

The development of novel therapeutics is a lengthy process that draws from fundamental research discoveries to inform downstream innovation and applied clinical work (1). FDA approval requires preclinical lab experiments in animal models and clinical trials and humans. Only after safety and efficacy are demonstrated in clinical work, and intellectual property established for novel innovations, is regulatory approval generally achieved. Researchers have estimated a time lag of 17 years between the basic research supporting the development of a new drug and the ultimate healthcare practice (2–4). In principle, targeted efforts by the scientific community and science funders could accelerate the discoveries that drive therapeutic development (5,6). In practice, it is challenging for scientists, funders, and pharmaceutical companies to identify the most promising research avenues at an early stage (7,8).

To overcome this lack of information at early stages, researchers have turned to machine learning models to identify promising research earlier than subject matter experts. Prior research has focused on making predictions about specific clinical trial outcomes based on the publicly available clinical trial data deposited at government repositories such as ClinicalTrials.gov. These draw on features such as study design (9), free text descriptions of the trial (10), and industry participation and enrollment criteria (11). Many models focus on individual trial performance (12), but others also predict eventual approval (10,13,14). Late-stage trials in particular convey significant predictive power on FDA approvals. This is because they are the most statistically powered and because their existence represents a large financial bet by a well-informed stakeholder that approval is likely. However, a major disadvantage of relying on late-stage trial data is that by the time a Phase 3 (or even Phase 2) clinical trial has been initiated, most of the decision-making opportunities that could accelerate therapeutic development have already passed.

We asked if it is possible to link predictions of FDA approval not to individual, trial-specific data, but to the scientific literature that preceded this work. Using this reasoning, it might be possible to link FDA approvals not just to their preceding trials, but also to the collective efforts of scientists that culminated in a novel therapy. To accomplish this, it is necessary to subdivide the literature into clusters that each represent the collective efforts of scientists in which pivotal clinical trials will eventually appear (15). Using publication clusters that can be tracked over time using historical data, it may be possible to use the emergence of one of these pivotal trials linked to an FDA approval as a positive signal for training machine learning models, where the features are the properties of the cluster as it existed prior to trial completion. These topic clusters could provide information about the character, growth, and reception of the scientific community for any given topic (16,17). The cumulative nature of drug development makes the cluster level approach particularly suitable for our context. In addition, prior models carry a significant limitation: most predictive power is carried by late stage trials, after which the key opportunities to change or accelerate research directions have already passed (10,13,14). Furthermore, missing data in clinical trials datasets are challenging for such modeling approaches (10). This may be particularly problematic at early clinical trial stages, before sponsors have uploaded complete trial results. Many sponsors have not completed depositing trial information by the time of FDA approval (18). By using a modeling approach that studies the prior biomedical literature, earlier predictions can in principle be made in a time frame more compatible with informing decision-making. Furthermore, this approach does not require any particular trials to already be underway, and inferences can be drawn from the research trajectory itself.

Here, we test the degree to which such Research Cluster (RC)-level information can provide insight into the likelihood that a given topic will receive a novel FDA-approved therapeutic in the future using machine learning. This approach complements, rather than replaces, models based on individual clinical trials. We leverage publication and citation dynamics among RCs in order to estimate the likelihood of a future FDA drug approval in a research topic, regardless of whether clinical trials have been initiated for a particular drug-disease pair. We find that the RC properties hold significant predictive power in identifying those topics most likely to yield an FDA approval, flagging RCs that yield novel drug approvals over 13 years in advance.

## Methods

IRB approval was not necessary because this study analyzed publicly available data.

### Biomedical literature citation data

To generate research clusters from publications, we used the 2021-06 NIH Open Citation Collection dataset consisting of over 640,000,000 direct citations and corresponding NIH iCite Database Snapshot (19–23). The Open Citation Collection was used because it is the most comprehensive citation dataset of biomedical citation linkages (20) and the best aligned with the biomedical focus of the present study. We cut the citation graph using the Leiden algorithm (15) to generate clusters of scientific publications that represent closely related topic clusters (Fig. 1).

**Fig. 1.**
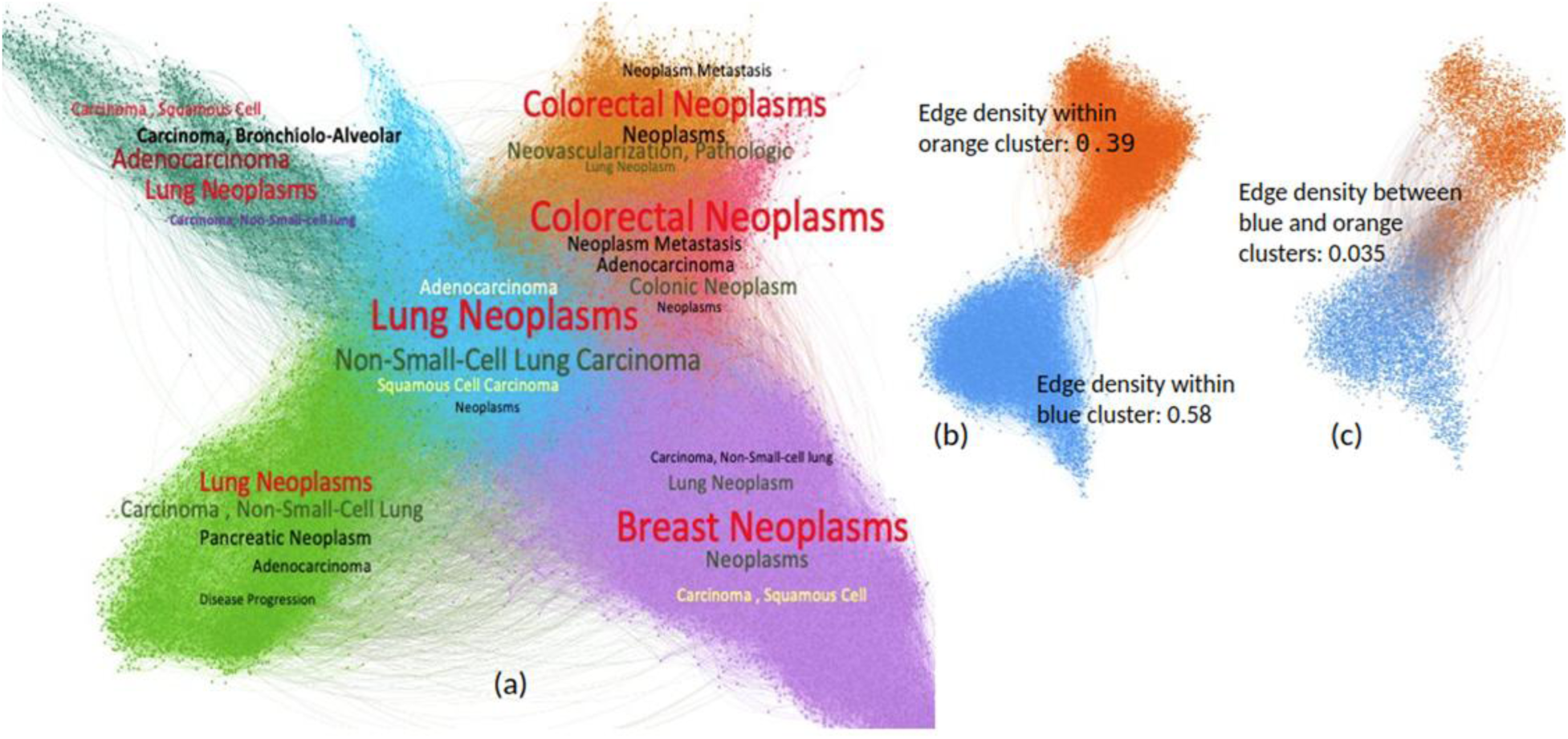
Topic clusters in “Alecensa” subnetwork and Intra-Inter cluster connectivity. (a) Subnetwork related to the research cluster of the drug “Alecensa” (blue) and five of its most interlinked neighbors by citation edges with the most frequent disease terms associated with the clusters. Colors represent different clusters where nodes are individual articles and citation between two articles denotes edge. (b, c) Two clusters from (a) with either between-cluster citation edges masked (b) or within-cluster edges masked (c). The articles (nodes) are the same in (b) and (c), only the edges are changed for visual effect. (b) Citation edges within the same clusters (both citing and cited articles are in the same cluster). Articles within the same cluster have a high density of citation links (edge density within the blue cluster is 0.58 and within the orange cluster is 0.39) (c) Citation edges between articles in two different clusters (citing and cited articles are in two different clusters). Articles of two closely related but different clusters have a comparatively lower density of citation links between them (edge density at the intersection of orange and blue clusters is 0.035).

### Graph Clustering and Research Cluster Formation

We used the graph clustering Leiden algorithm (15) to form RCs derived from the NIH Open Citation Collection citation graph (21,22). Because clustering is performed on article-level PubMed IDs (24) rather than author names, the known author disambiguation limitations of the NIH Open Citation Collection do not affect our analysis. While the NIH Open Citation Collection has known limitations linked to coverage of PubMed (in non-English language publications), it is the most comprehensive citation graph of the biomedical literature (20,25). The Leiden algorithm generates internally connected communities where communities are locally optimally assigned, and it outperforms the baseline Louvain algorithm in terms of speed and better partitioning. The algorithm clusters highly cohesive and semantically aligned articles within the same cluster (Supplementary Fig. 1). As we aimed to predict novel therapeutics from biomedical literature, it was important to operationalize this concept. RCs are composed of clusters of scientific articles labeled with their PubMed Identifiers (PMIDs). However, there is no structured data in PubMed about which papers supported the approval of a novel therapeutic. To map this, we leveraged datasets linking specific FDA approvals of New Drug Applications back to their supporting clinical trials registered in ClinicalTrials.Gov. We map the clinical trials to the associated publications through ClinicalTrials.Gov as it includes the publications associated with the trial. Not all clinical trials are pivotal to the FDA’s decision-making, however, and we wanted to ensure that the RCs we labeled as ‘positive’ were linked to pivotal clinical trials. We cross-validated each ClinicalTrials.Gov identifier (abbreviated as NCT ID) with public documentation from the FDA indicating for each New Drug Application (NDA) approval which trials were considered pivotal for the FDA’s decision-making. RCs containing the PMID of a pivotal trial were used as a starting point for identifying positive RCs. Because each set of RCs is generated from a snapshot of the citation graph at a single point in time, we generated historical citation graphs that would have been available at the end of each publication year. We tracked RCs over time as they evolved, grew, and split into smaller, novel RCs (i.e. newly differentiated topic clusters that emerge as spin-offs of a parent RC) that represent new topics of scientific inquiry.

Graph clustering uses edges (e.g. citations) as the substrate for assigning nodes (e.g. papers) to a cluster. Previous results have shown that pruning a citation graph to remove certain types of local network structures can improve analytical performance (26–29). We asked whether pruned networks preserving different types of local network structures could improve modeling performance. Before engaging in feature generation, selection, and predictive modeling, we first generated four citation graphs from which to generate clusters that could be used for predictive modeling (Supplementary Fig 2), based on edge types that have demonstrated predictive value in the past (26).

The first was unmodified direct citations. The second was only citation edges where the cited document was co-cited by other articles from the citing article’s reference list (“co-cited by reference”, (26)). This type of edge could be interpreted as a confirmation from the other related documents in the reference list that the given cited article is relevant or authoritative. The third edge type we tested was a citation that represented both a direct citation and a co-citation relationship (“direct-and-co-citation”). This local network structure occurs when both papers in a citing-referenced article pair are cited in a subsequent study, possibly confirming their mutual relevance to subsequent work. The fourth edge type (“trimmed”) we tested was the opposite of the “co-cited by reference” set and is conceptually similar to the longest-path citation relationship described by (29,30). This represents the trimmed citation network that removes any citation that was also recapitulated by the “co-cited by reference” set. It does not strictly speaking represent the longest path between two nodes but removes redundancy in the citation network.

### Features

To generate features that might hold predictive power, we relied on metadata disseminated through the biomedical research databases PubMed and iCite (Table 1). In particular, we included summary statistics about a cluster’s scientific influence (22), the human-, animal-, and molecular/cellular domain of the constituent research articles (21,31), non-research content of a cluster (i.e. editorial articles and reviews). We also included flocking behavior by scientists into a scientific topic (e.g. new papers published within the previous 2-3 years) and the cohesion of a subject over time (see Supplementary Fig 3). Note that RC-level growth features aggregate contributions from many research groups. An RC typically contains hundreds to thousands of papers from many institutions, so the growth signal reflects broad community interest rather than the output of any single laboratory. Finally, we included features such as reference age (the number of years between publications from a cluster and the works that they cited), the fraction of papers funded by the NIH and the proportion of papers in a cluster that constitute clinical citations.

**Table 1.** Features considered for training.

|  |  |
| --- | --- |
| Recent growth (0-1 and 1-2) | These features measure the recent explosion in the number of research focused on a particular topic and in the particular cluster. We measure the new papers published within two 2-year sliding windows, the previous 0-1 years, and 1-2 years. The percent of papers published in the previous 0–1 years and 1–2 years. |
| RCR of the top 95th percentile paper in the cluster | This feature measures the intellectual influence of the articles within a particular cluster. The influence is measured by the metric RCR (22). |
| Growth of second biggest ancestor | Proportion of papers in the second biggest ancestor cluster (previous year's cluster with highest level of PMID overlap to the present year's cluster) of the terminal year that published in the past 0-1 years. |
| Human score | The occurrence/avg of human focused research within the cluster. Defined in detail at (21) |
| Median RCR of the papers in the cluster | The median RCR of the papers within the cluster (22). |
| Animal score | The occurrence/avg of animal focused research within the cluster. Defined in detail at (21). |
| Molecular / cellular score | The occurrence/avg of molecular-cellular focused research within the cluster. Defined in detail at (21). |
| Clinical score | The proportion of clinical articles in the cluster. Defined in detail at (21). |
| Primary research score | The proportion of research articles in the cluster, as opposed to secondary articles like reviews, or non-peer-reviewed items like preprints (34,35). Having higher proportion of research articles in a cluster is negatively correlated to drug intervention. |
| Clinical citation | Proportion of articles cited by clinical articles (21). Having clinical citation indicates the potential of the research topic to translate into clinical research |
| Percent of papers funded by NIH | The fraction of papers funded by the NIH. |
| Share from the biggest ancestor | Proportion of the articles that are from the biggest ancestor in the terminal year<br>Indicates stability of clusters over time. |
| Share from top 5 ancestor | Proportion of the articles that are coming from the top five ancestors in the terminal year. Indicates stability of clusters over time. |
| Reference age | The number of years between publications from a cluster and the works that they cited. |
| Weighted RCR | Sum of article RCRs in that cluster (22). |
| Size of previous year's closest ancestor cluster | No. of papers in the biggest ancestor cluster of the terminal year. |
| Share from the second biggest ancestor | Proportion of paper coming from the biggest ancestor cluster of the terminal year. |
| Size of previous year's second closest ancestor cluster | No. of papers in the second biggest ancestor of the terminal year. |
| Share from the second biggest ancestor | Proportion of paper coming from the second biggest ancestor of the terminal year. |

We generated feature sets that fall into five broad categories based on the type of information they provide (Table 1). The first category measures the entrance of new papers into a topic as represented by an RC; this measurable flocking behavior by scientists into a topic is a strong indicator that it is scientifically exciting, represented by recent growth. The second set utilizes the National Library of Medicine’s Medical Subject Heading terms and their use in generating a translational space termed the Triangle of Biomedicine (21,31), the latter of which can help to predict future early-stage bench-to-bedside activity (21). The third set directly measures clinical publication and citation activity, for which there is now comprehensive structured data available (19,20,23). The fourth set examines measures of scientific influence and productivity (22,32,33). The last set of features measures the dynamism of an RC. Each RC is a cluster of papers linked to a given snapshot in time, so for every new publication year, the cognate RC is somewhat different, acquiring new papers and possibly spawning new RCs. Together, these features provide significant diagnostic signals to the machine learning model to make accurate predictions about individual RCs’ drug development likelihood.

The detailed definitions of the features that we considered are described below. Full feature correlations with the outcome variable are shown in Supplementary Fig. 4.

### FDA Approval Dataset

In order to build a data pipeline to generate training and test datasets, clinical trial and publication data associated with each drug were aggregated. We leveraged earlier work that disseminated clinical trial data assembled by the Good Pharma Scorecard organization, linking approvals to ClinicalTrials.gov and PubMed (18,36–40). These curated datasets did not always distinguish between pivotal trials used in FDA decision-making and those that were not. Starting from these, we curated seminal clinical trials that the FDA documented were directly used in the agency’s decision-making to conduct our analysis. Pivotal trials were curated from FDA documents and publications associated with each of the drugs were linked so that these could be matched to publication clusters. Clusters that contained publications describing clinical trial results designated as pivotal in FDA documents were labeled as positive, and others as negative.

Drug-related data were integrated from publicly available clinical trial datasets, trial registries including ClinicalTrials.gov, corporate registries, and the WHO’s International Clinical Trials Registry Platform; PubMed; and linked data repositories (clinicalstudydatarequest.com and yoda.yale.edu). These included new therapeutic biologics, molecular entities, and drugs approved by the FDA in the years 2012, 2014, 2015, 2016, and 2017 from Drugs@FDA. For years 2012, 2014, and 2016, these only included new drug applications and biologics sponsored by the 20 largest companies based on the particular year’s market capitalization ranking.

### Model Training and Validation

We formulated our problem as a supervised binary classification and built a machine-learning model to predict the success or failure of a research cluster in generating drug intervention as described above. We used a balanced number of positive and negative RC samples from different years for preliminary testing and feature selection. We built several classifiers including regularized maximum likelihood logistic regression modeling, random forests, extreme gradient boosting, support vector machine, and naive bayes models, and finally selected the maximum likelihood logistic regression model as it outperformed the others. We identified the most important features of the RCs from each of the four edge types. The importance of the most predictive features and their correlation with the outcome label is shown in the supplement (Supplementary Figs 3 and 4). The top five features were the fraction of papers in an RC that are recent (either published in the most recent 0-1 years or 1-2 years), the fraction of papers that are primary research (negatively correlated, e.g. more editorials and reviews are a positive indicator of likely future therapeutic breakthroughs), the fraction of articles that are clinical, and the average Human score of papers in that RC.

To maximize model accuracy, we performed feature selection independently for each of the four edge types and trained four different models based on these features for the four edge types. We tested performance using 5,10, and 20-fold cross-validated F1 scores. Table 2 shows these results. Although we used a stratified model for training, Supplementary Table 2 and Supplementary Fig. 5 show results on a full selection of research clusters. The performance significantly varies across RCs generated using different edge types. The RCs using the “trimmed” edge type significantly outperformed the other edge types. We built a single model to compare the performance of all edge types using a common set of features which we refer to as the “unified” model in the rest of the paper. It yielded similar results on per year analysis (Supplementary Table 2), confirming the outperformance of the trimmed edge type again. As the trimmed edge type outperformed the other edge types in the unified model, we conducted our subsequent time-series validation analysis on the unified model with the trimmed edge type (Supplementary Fig 5).

**Table 2:** Cross-validated F1 score of the best models for four edge types.

| Edge type | 5-fold cross-validated F1 score | 10-fold cross-validated F1 score | 20-fold cross-validated F1 score |
| --- | --- | --- | --- |
| Trimmed | 0.8325 | 0.8421 | 0.8404 |
| Direct citation | 0.7441 | 0.7554 | 0.7806 |
| Co-cited by reference | 0.7619 | 0.7861 | 0.8039 |
| Direct- and co-citation | 0.7519 | 0.7488 | 0.7802 |

### Pivotal Trials and Associated Publications

For each drug in the combined data, we extracted the approval letter, summary review, medical review, chemistry review, pharmacology review, statistical review, and other clinical and non-clinical review reports from the FDA approval package (Fig 2). We extracted the clinical trials listed in medical review and approval letters and extracted the trial identifiers from there and matched them with the ClinicalTrials.gov website using the identifier. We matched the other Study ID Numbers, titles, number of participants, and phase number reported in the review letter and the official NCT registration website for the particular trial with the review letter so that we can be confident that we found the correct trial that the approval review letter referred to. In cases where a medical review was not available or did not report the associated trials, we used the summary review, chemistry review, and clinical review to find the trial information. We identified trials as pivotal if the trials were explicitly mentioned as pivotal or primary or key in the review letters. From the trial registration website of each trial ID, we found the relevant PubMed IDs associated with the trial. If multiple articles were indexed to a particular trial id, we manually checked the abstract of the publications and filtered the articles that were actually reporting the result and reported the same trial registration id. We also matched the number of participants, and the phase of trials reported in the ClinicalTrials.gov website and the associated publications. Most trials mapped to one cluster trajectory, but if not, then the research cluster was randomly selected such that each drug was matched to only one cluster.

**Fig. 2.**
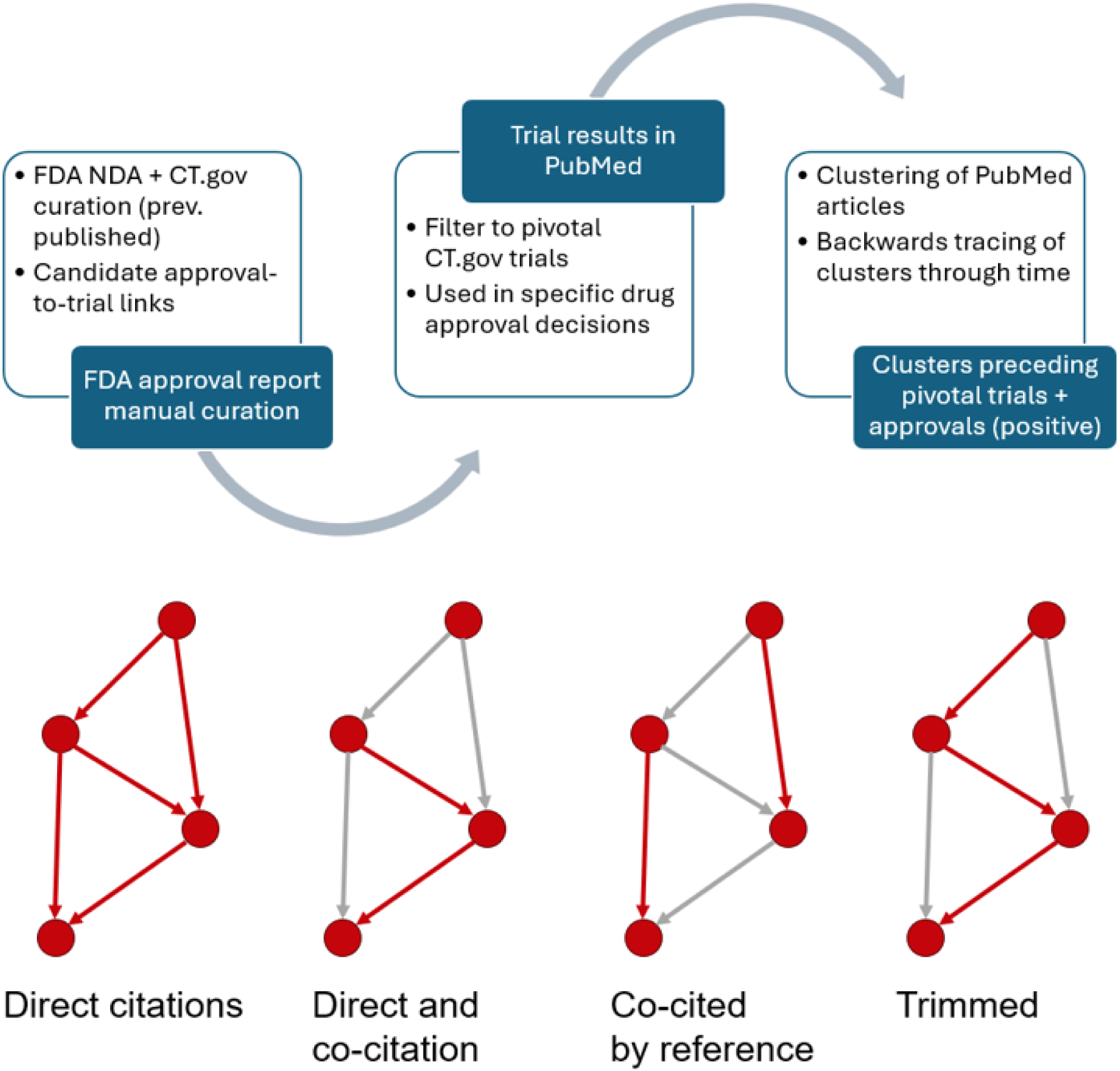
Prediction workflow for this study. (Top) Curation process for identifying positive clusters, starting with previously published candidate FDA NDA approval-to-trial linkages (18,36–40). (Bottom) Edge types tested, each of which correspond to a different type of local network structure (also see Methods, Graph Clustering and Research Cluster Formation).

Because positive clusters are identified through the appearance of a pivotal trial’s PubMed ID (linked to its ClinicalTrials.gov NCT ID and FDA NDA approval), drugs are not searched in the graph by name. The clusters measure topic level properties like recent growth, clinical orientation, and influence, rather than drug-specific content. This is why the model is able to learn features predictive of approvals despite lacking a specific view on drug properties.

### Statistics and Reproducibility

Model accuracy measures were calculated using Scikit-Learn’s F1-score, precision, and recall. Paired t-testing (two-sided) on disease-matched clusters was performed in R using research clusters matched to exemplar clusters on exact matches to historical year and top disease MeSH term, and matched to the nearest cluster size after applying those two exact matching criteria, using the data available in the Data and Code Availability section (41). Fisher’s exact testing (two-sided) was done in R with the exact counts described in the Results, Failed Drugs section.

## Results

We began by generating clusters of the scientific literature in which FDA New Drug Application (NDA) approvals could be linked through their approval documents, clinical trial registries, and subsequent publications (see Methods). We used the Leiden clustering algorithm to subdivide the entire biomedical citation graph represented by the National Institutes of Health (NIH) Open Citation Collection (19), generating RCs for each historical year using only citation data that would have been contemporaneously available at the time. Drug approval data spanning 2012-2017 was linked to RCs by when an article reporting the results of a pivotal clinical trial appeared in that cluster.

Fig. 1 shows a small proportion of the direct citation network related to the research of the drug “Alecensa” (trade name) or “Alectinib” (nonproprietary name). Alecensa is a drug that is used to treat non-small-cell lung cancer and was approved by the FDA in 2015. Note that this particular subnetwork was chosen for illustration; the full model generates tens of thousands of such RCs. The center blue RC is associated with a pivotal Alecensa clinical trial that supported its FDA approval. The neighboring RCs are those that are most highly connected to the center cluster by citation edges. The disease terms associated with each cluster in Fig. 1(a) show that the clustering algorithm accurately separates clusters based on their semantics and contents and places RCs focusing on similar diseases closer in terms of citation edges. Fig. 1(b) shows the citation edges between papers within the same cluster. Because these papers cite each other frequently, these papers are grouped together by the graph clustering algorithm. Fig. 1(c) shows the between-cluster citation edges, which are comparatively sparser, which leads to cuts by the clustering algorithm that separates these respective groups of papers into two different clusters. These figures indicate that the topical cohesion and individuality of different scientific fields are well-represented by the graph clustering approach. This is confirmed by semantic similarity analysis of the papers within and between separate topic clusters. Semantic cosine similarity is much higher for within-cluster article pairs than for between-cluster article pairs (Supplement Fig 1).

We asked whether this approach can be used to predict in advance the emergence of novel therapeutics. In order to operationalize this question, it is necessary to have a defined outcome measure, as well as features in which a machine learning system can detect predictive patterns. As an outcome measure, we leveraged prior work (18,36–40) that identified pivotal clinical trials that supported the approval of new drugs from 2012-2017 (see Methods, FDA Approval Dataset and Fig. 2). These take the structure of approved drug names, linked to the ClinicalTrials.gov NCT identifiers, which are then linked to the scientific publications in PubMed describing the results of these trials. Because unsupervised topic clusters take the form of lists of PubMed IDs (PMIDs), we used the presence of at least one such pivotal trial paper to label a cluster as positive, and their absence to label clusters as negative.

For generating features on which a machine learning algorithm can learn, we used historical snapshots of RCs truncated to only contain data as they would have appeared in each historical year back to 1980. RCs were linked to those of prior years by measuring the number of overlapping papers from year to year; the prior RC with the most overlapping papers was linked to the following year’s RC in order to trace the development of a topic over time. Features can be measured at multiple levels (see Methods, Features section): RC level (e.g. cluster growth, age of papers in the cluster, cluster stability), or aggregates of article level data (e.g. percent human, animal, or molecular/cellular biology papers, percent that have been cited by a clinical study, percent that were NIH-funded, or article influence measured by the Relative Citation Ratio). Our feature selection (Methods and Supplementary Methods, Feature Selection) determined that thirteen of these features were the most salient for predicting which RCs would host a pivotal trial’s paper in the coming years. Features after selection are shown in Supplementary Figs 3 and 4. Measures of recent growth were positively associated with therapeutic approval. This may represent a measurable flocking of researchers into an area anticipated to yield exciting results in the future. An above-average fraction of primary research articles was a strong negative predictor. RCs leading to a novel drug approval are instead enriched in reviews and news articles, reducing the share of primary research papers. Historical article influence measured by time series snapshots of the Relative Citation Ratio and share of clinical articles were both positively associated with drug approval. On the stratified sample used for training, these features achieved an F1 score of 0.78 in 20-fold cross-validation on the direct citation network. This demonstrates that future drug approval can be predicted based on a combination of the RC dynamics and their article properties in prior years leading up to approval.

In our prior work, we noted that local citation information carries predictive power in predicting whether a given publication-to-publication citation is substantive in nature (26). Because our unsupervised clustering approach uses citations as its substrate, we decided to test whether particular types of local citation network structure can convey predictive power, before the cluster features are even generated. For this reason, we tested the predictive power of our selected features on RCs generated on four types of citation edges (Fig. 2). The trimmed network structure yielded the best performance of the four by a large margin, achieving an F1 score above 0.84 in cross-validation (Table 2). Removing the common knowledge transmission pathways as represented by overlapping citation links among the papers in an article’s reference list, and restricting to only those that represent novel information transmission pathways in this subset, carries substantially more predictive power than leaving the original citation network intact.

### Time Series Validation

To validate predictive accuracy and test how early our model can identify drug-related research clusters, we identified the trajectories (chains of RCs from year to year containing the pivotal trials supporting FDA approval) of 99 individual exemplar drugs. For each drug, we generate the trajectory of research clusters associated with the drug by tracing the clusters associated with the pivotal publications of the relevant clinical trials over several years. We generated a test dataset with approximately 370 data points corresponding to the publications of pivotal trials of those drugs. First, we analyzed the distribution of prediction scores for exemplar drug clusters on different edge types using our unified model (Supplement Fig. 6). The “trimmed” edge type showed the greatest degree of discrimination between positive (exemplar) clusters vs. negatives and outperformed the other three edge types in detecting positive exemplar drug clusters (Supplementary Table 3).

Predictive analytics should yield timely signals that can predict, years in advance, the outcome of interest. To determine how far in advance our model signals the likelihood of a future approved drug, we examined the distribution of prediction scores relative to the publication of pivotal trials and FDA approval dates. We focused on the “trimmed” edge type as it outperformed other edges by a large margin. For a particular exemplar drug, we selected the RCs prior to the publication of results. In Fig. 3, we compare the prediction score given by the unified model for the exemplar drug clusters to randomly selected negative clusters. We quantify prediction scores relative to the FDA approval year. For comparison, we randomly selected a similar number of non-exemplar data points (Fig. 3b and 3d). Fig. 3a and 3c show that the prediction scores of RCs in the research trajectory of an exemplar cluster are much higher than a randomly sampled set of RCs. Fig. 3c shows that high prediction scores precede FDA approval many years in advance in contrast to randomly sampled (presumably negative) RCs. These large prediction score differences demonstrate that it is possible to identify a novel FDA approval many years in advance. The timeline distribution of high prediction scores for the 99 exemplar drugs shows that almost 80 percent of drug clusters provide a signal before their approval with more than 65 percent providing signals at least 8 years before their approval years (Fig 4).

**Fig 3.**
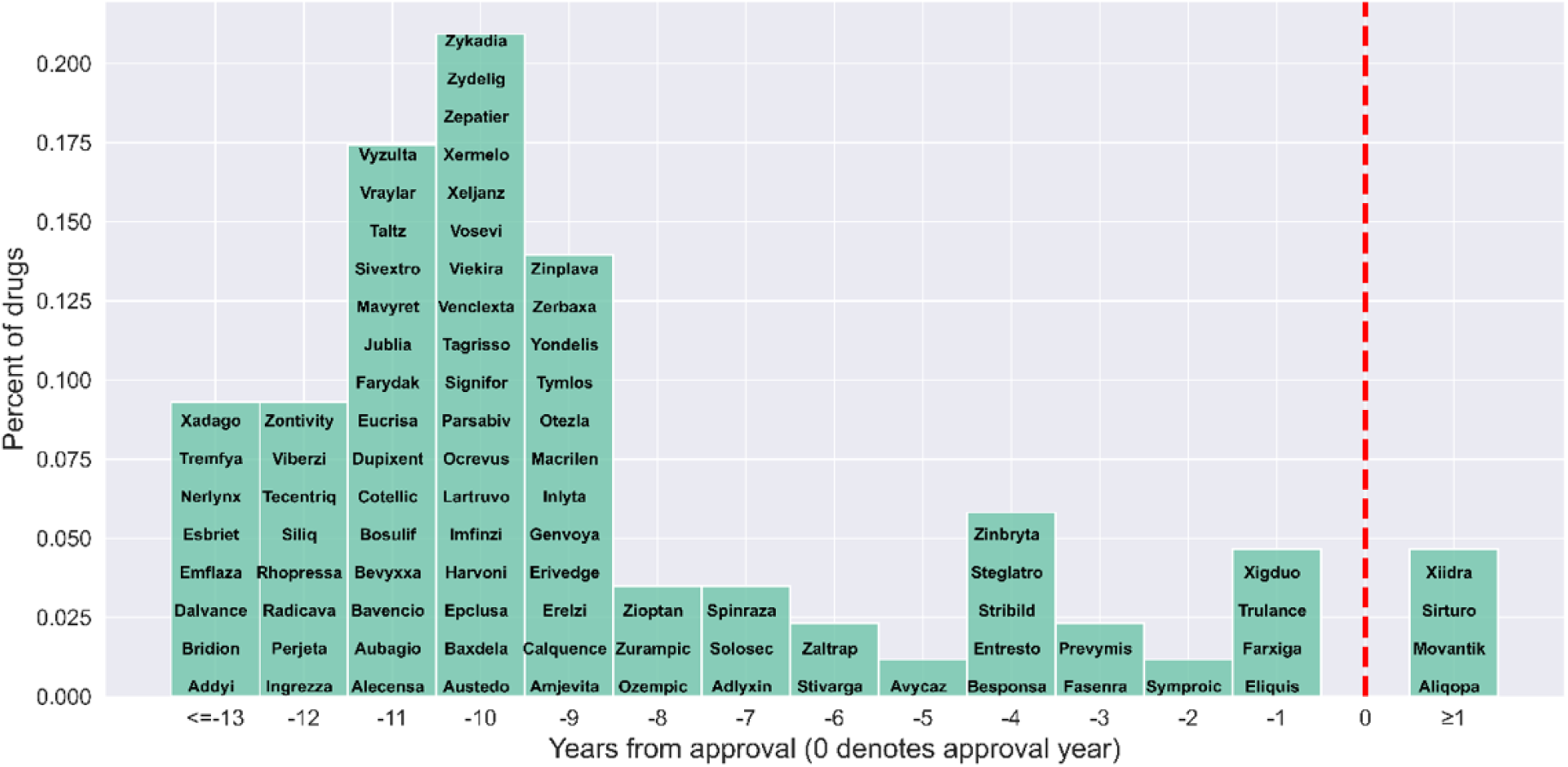
Prediction score on exemplar and random non-exemplar clusters. Year 0 denotes the approval year of the drugs and the negative years denote the years before approval. The red dotted line is the prediction score threshold for denoting positive prediction (see Supplementary Table 1). (a) More data points are above the threshold for exemplar clusters, their ancestors, and their descendants and the fitted regression line consistently remains at or above the threshold line. (b) More data points are at the lower half of the prediction score for random clusters, their ancestors, and descendants. The fitted regression line (bands, 95% confidence intervals) is much lower than the exemplar clusters and the skew of distribution is opposite to the exemplar clusters in (a). (c) Box plot of the prediction made by the model on the exemplar drug clusters, their ancestors, and their descendants (n = 906 clusters). The higher prediction score starts to appear several years before the approval year 0. (d) Box plot of the prediction made by the model on the random clusters, their ancestors, and their descendants (n = 845 clusters). The range of prediction scores for different years is much lower compared to the exemplar clusters in (c). For box and whisker plots: center line, median; box limits, upper and lower quartiles; whiskers, 1.5x interquartile range; points, outliers.

**Fig 4:**
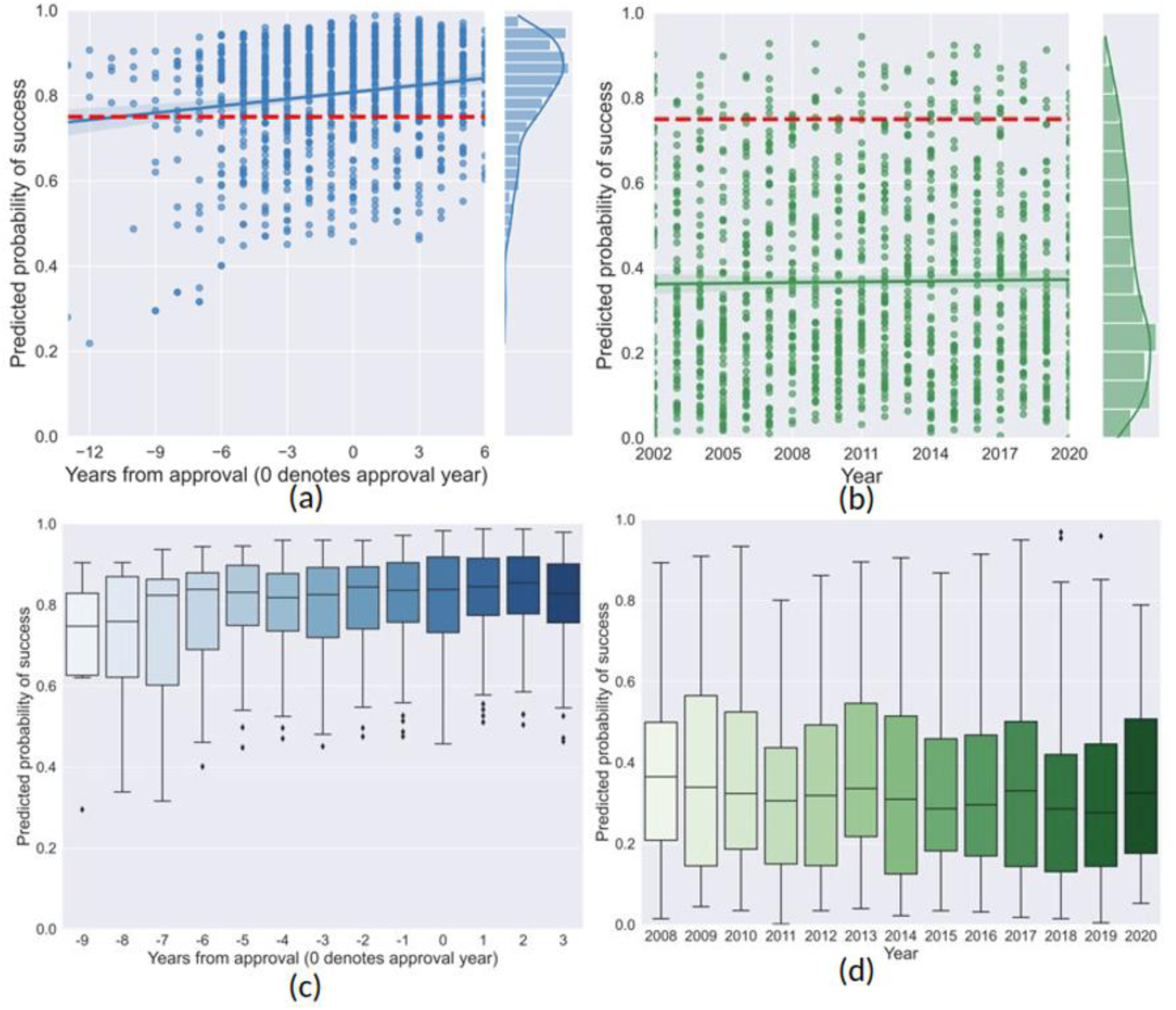
Exemplar drug clusters identification timeline. The histogram shows the timeline of the 99 exemplar drugs when our model provides the first positive signal (high prediction score) for the specific drugs. **0 is the approval year.**

### Out-of-Sample Blockbuster Drugs

Out of necessity, we relied on curated datasets linking FDA approvals to pivotal clinical trials and their associated publications. However, these data represent only a small minority of the true positive trials and associated RCs, in large part because prior to the Food and Drug Administration Amendments Act (FDAAA) of 2007, organizations were not required to deposit such data in public domain repositories. In addition, our training dataset relied primarily on the FDA Orange Book documenting small-molecule drugs but potentially overlooking macromolecule treatments documented in the FDA Purple Book. As a result, “false positives” (Supplementary Table 2) in our dataset may represent true-positive RCs that were simply out of scope for generating our training data. To test this hypothesis, we analyzed prediction scores for RCs of out-of-sample blockbuster drugs (identified via PharmaCompass). For each drug, we extracted the research cluster of the associated publications across several years and made contemporaneous predictions (e.g. predictions that used information available at that historical time) about different data points along the trajectory before and after the approval.

For 8 out of 10 drugs, our model generates scores exceeding our previously established threshold (>=0.75) before their respective approval years. The model is able to identify 7 out of the 10 drug-related clusters at least 2 years before their approval. 8 out of 10 drug trajectories first exceeded the prediction threshold before approval, and this high score was sustained for several subsequent years (Fig. 5). Notably, 6 out of 10 drug trajectories appeared to be promising for the first time several years prior to approval and maintained the higher score for several years ahead, confirming the results of our analysis of in-sample exemplar drugs. This result confirms the generalizability of our modeling approach even to drug types that were out-of-scope for the training of the model.

**Fig 5.**
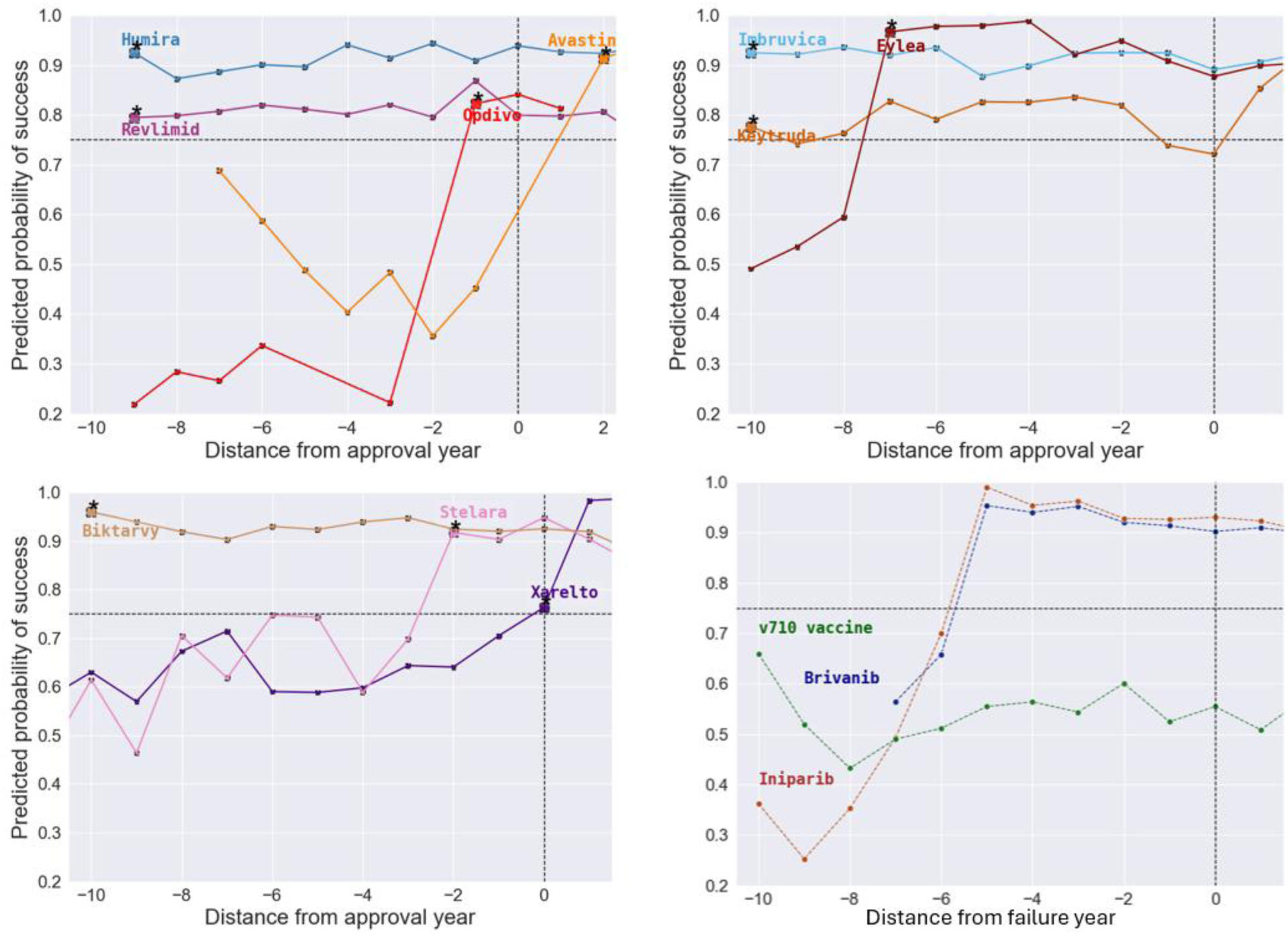
Prediction score across the trajectories of 10 out-of-sample blockbuster drugs and near-misses. The “*” denotes the first appearance of higher prediction (>0.75) in the development trajectory of the drug and year “0” in x-axis denotes the approval year (top, and lower left). The horizontal line denoted the prediction score cutoff for higher prediction and the vertical line denotes the approval year. 8 out of 10 drugs yielded higher prediction scores before their approval years and 6 of those (trade names - Humira, Revlimid, Eylea, Imbruvica, Keytruda, Biktarvy) appeared to be promising at least 7 years before their approval. A selection of failed drug approvals that made it to large-scale phase 3 trials but had divergent results is shown on the lower right panel. These RCs also typically showed high prediction scores.

### Prediction Signals Appear Before Phase 2 Trial Start Dates

In order to conceptually anchor this finding, we compared the year of the earliest signal against the start dates of the pivotal phase 2 trials that were cited in the FDA’s decision-making, and which were available on ClinicalTrials.gov. Not all drugs have a phase 2 trial designated as pivotal in the FDA’s public reports, and some of these may predate the 2008 public reporting requirements on ClinicalTrials.gov. However, for 29 drugs, we were able to link a pivotal phase 2 trial to a clinical trial registration, and found our prediction signals predated these phase 2 trial start dates 86% of the time. This means that our prediction signals often appear before the corresponding phase 2 trials have even begun.

### Failed Drugs

In order to further test our modeling process, we examined a set of 21 drugs that, while promising, had divergent phase 2 and phase 3 trial results, leading to their failure (42). One drug was excluded from analysis because it was approved, meeting our positive criterion, and later withdrawn. We examined the prediction scores of their linked RCs leading up to the time the failure was known. 13 of these 21 drugs had a positive signal prior to the known failure, on average 7 years ahead of the failure. This is not statistically significantly different from the 81 of 99 in our main sample (p = 0.55, Fisher’s exact test, n = 120, t = -15.866). Examples of three failed drugs are shown in Fig. 5. After rising to the prediction threshold, scores typically remained high for these failed drugs.

Based on these results, combined with our analysis of phase 2 trial start dates, we interpret these as an indicator that the modeling process described here is sending early signals that a topic looks very therapeutically promising, rather than a particular drug will successfully transition from phase 2 to phase 3 trials. For example, while iniparib failed in these trials, other drugs in the same class for the same indication such as olaparib and niraparib succeeded in the subsequent years. The predictions generated here may indicate that a line of research is promising, not that individual applications are likely to succeed.

### Strengths and Weaknesses of Modeling the Scientific Literature

Previous predictive approaches have sought to model the likelihood of a drug’s success at achieving FDA approval based on the linked documents uploaded to ClinicalTrials.Gov (10,12). However, our approach is not drug specific. Rather, it is scientific topic-specific, and we asked whether our approach performed better at predicting novel therapeutics at the level of drugs vs. diseases.

We analyzed the MeSH (Medical Subject Headings) terms associated with RCs to quantify the content of the cluster. Interestingly, our analysis suggests that the disease that a drug is primarily intended to treat appears in top MeSH terms consistently compared to the rest of the terms in those clusters (Fig. 6). However, our analysis of drug-specific terms suggests that our modeling approach does not perform well at predicting the specific drug that is likely to be approved. In contrast to this lack of specificity with respect to future approved drugs, the diseases that are likely to be treated are easily discerned, demonstrating that the model reliably identifies disease areas likely to be of interest to patient groups and the public.

**Fig 6.**
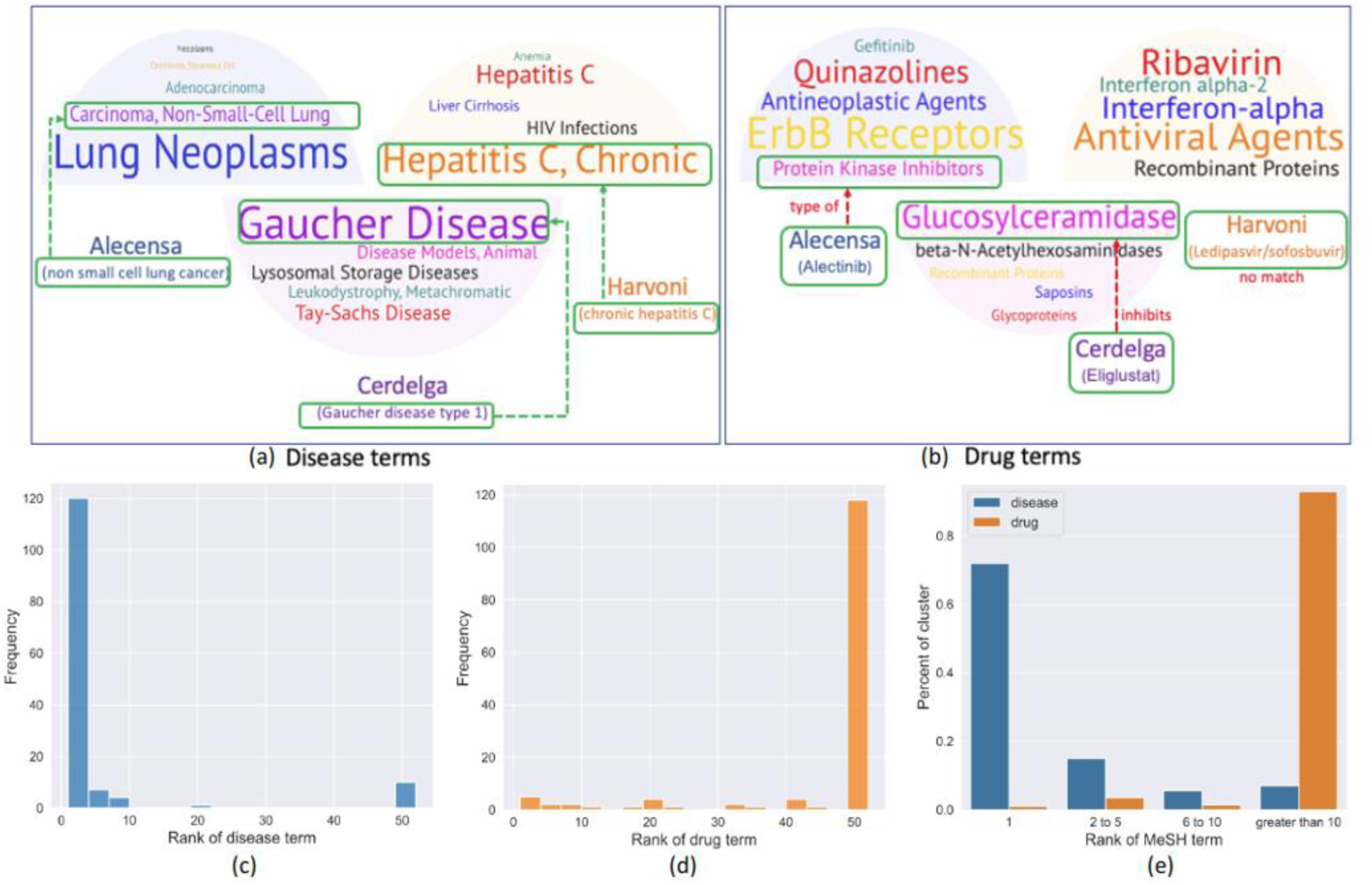
Drug and Disease MeSH term distribution of the exemplar drug. (a-b): Most frequent disease and drug MeSH terms in the research clusters of three exemplar drugs - Alecensa, Harvoni, and Cerdelga. (a) Disease terms are well representative of the actual disease name associated with the drugs(Harvoni: chronic hepatitis C), Alecensa (non-small cell lung cancer), Cerdelga (Gaucher disease type 1)). (b) Drug terms are scattered and vague and less representative of the actual drug name and synonyms. (c-e): Ranking distribution of the actual disease (c) and drug (d) names of the exemplar drugs appearing in the most frequent MeSH terms associated with their clusters. The diseases treated by the drugs in the clusters are consistently represented in the top disease MeSH terms associated with the clusters whereas the drug terms are not well represented in the top drug MeSH terms (e). Bin 50 contains all the clusters where the disease term does not appear in the top 50 terms.

One reason for this lack of drug specificity may be due to the nature of the RCs themselves and the timeline of the signals that are measured. RCs are typically several hundred to thousands of papers large, and therefore encompass the collective effort of whole scientific communities rather than the output of individual labs or institutions. Thus, these may indicate the convergence of a pressing therapeutic need combined with an opportunity that is sensed by the participants in the research enterprise. As such, it might be clearer that a therapeutic breakthrough is suddenly possible, even in the absence of information of which particular approach will prove the most effective. This is consistent with the timeline of signals that we find, a mode of 10 years before approval, at which point safety testing has not yet been conducted. This combination of properties may be why diseases and not drugs are most effectively predicted.

### Disease Matched Clusters

In order to test whether our modeling approach is entirely disease-specific, we asked whether in general, clusters with specific disease MeSH terms receive positive labeling. We matched exemplar clusters to other clusters with the same top disease MeSH term from the same year, and the closest cluster size. Disease-matched clusters received prediction scores that were on average much lower than the exemplar clusters (Fig. 7). This was significantly lower than the exemplar clusters themselves (p < 2.2e-16, two-tailed paired t-test, n = 130, mead difference = 0.310, 95% confidence interval 0.271-0.349, t = 15.866, df = 129). Thus, while disease-matched clusters have a higher prediction score on average than a random negative sample (Fig. 2), focus on a particular disease does not guarantee a high prediction score. Feature importance scores (see Supplementary Methods) reinforce this finding: recent RC growth, high presence of secondary articles like reviews, and high RCR scores are among the most predictive features, and will not be uniformly high in all topics that focus on a particular disease.

**Fig 7.**
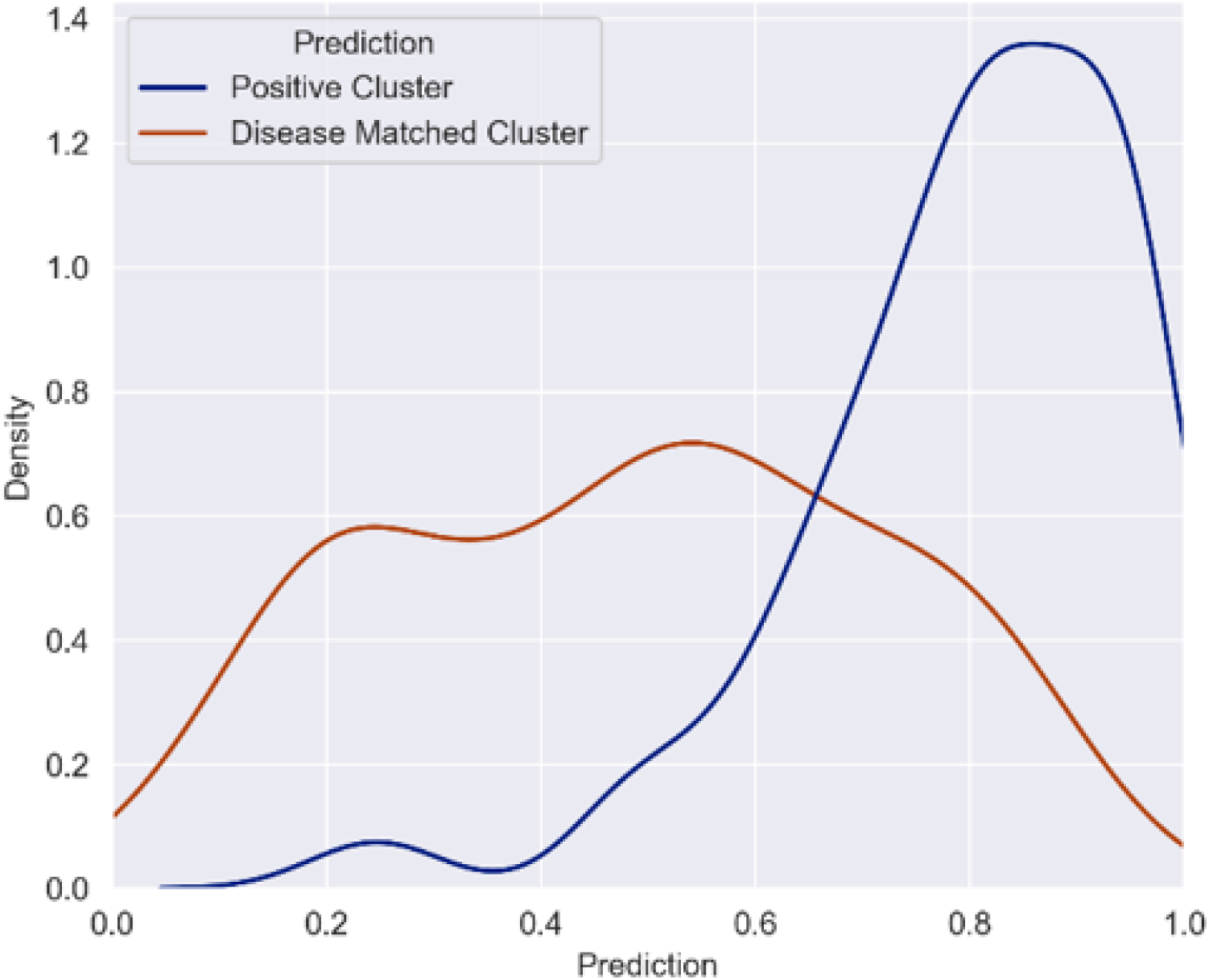
Prediction scores for positive clusters vs. those matched for the same disease (also matched for year and RC size). Studying the same disease is not sufficient to raise an RCs prediction score; instead other features such as recent explosive growth or high-influence articles may signal that a particular scientific topic is promising for novel therapeutic development (see Supplementary Fig. 4)

### Forward-Looking Predictions

To forecast the clusters most likely to yield novel therapeutics in the near future, we present 43 of the highest scoring RCs in Supplementary Table 4 based on a 2025 snapshot of the citation data. These range in size from cluster 0 (clusters are numbered in descending order by size, where 0 is the largest) focused on dysbiosis, to cluster 10984 focused on cross-infection. Consistent with our observations of stable prediction scores preceding therapeutic approval, we selected those clusters with scores > 0.75 for 2023-2025. We then mapped the most frequently occurring MeSH term onto these clusters and deduplicated on top MeSH term such that the highest scoring cluster for that disease is shown. The most frequently occurring MeSH term is mapped onto each cluster and used for deduplication; because this term tends toward broad disease categories and may not capture the nuances of each research trajectory, full cluster-level data are available at the link in our Data Availability section. These 43 clusters represent a small fraction of the tens of thousands of RCs in the 2025 snapshot, reflecting the model’s selectivity when applied to the comprehensive citation graph.

## Discussion

We find that research clusters signal information about which disease topics are the most likely to yield therapeutic breakthroughs. Future therapeutics are signaled by features including the sudden growth of a research cluster, large numbers of secondary research articles such as reviews, high article influence, and high numbers of clinical citations to papers in the cluster. This work provides a computational framework for predicting drug approval likelihood in any given scientific topic by combining citation knowledge networks, graph-clustering algorithms, and machine learning. Because many of these signals appear 13+ years in advance, this can provide a timely signal to researchers, institutions, and funders that a topic area is exceedingly promising for attention and resources.

Our modeling approach complements others that have focused on predicting whether a given clinical trial is likely to yield an FDA approval. Our approach does not incorporate specific domain information from posted clinical trials, and may in the future benefit from that inclusion. However, we sought to instead identify subfields that are among the most promising many years in advance. Since our signals reliably appeared before phase 2 trials have begun, they circumvent a main limitation of other modeling approaches: by the time the most accurate clinical trial-based prediction models can generate reliable signals from phase 3 trials, the opportunities to influence decision-making have passed. Our modeling approach does not reach as far back as the full 17 years that it is estimated bench-to-bedside translation takes, but it does significantly approach that time period. The modal signal here appeared 10 years prior to approval. This represents significant lead time for influencing the future direction of research investments.

Our approach has important limitations and opportunities for future improvement. As mentioned, we do not incorporate clinical trial information associated with the drugs into our model even though trial information may provide valuable insight into the drug approval. This represents an avenue for improving future predictive performance. The present work focuses on the biomedical research literature, but these types of approaches could be broadly employed across different knowledge graphs and outcome measures. Indeed, our companion paper (43) uses similar methodology uses similar methodology to predict, on a similar timeframe, when and where transformative breakthroughs of the type that win Lasker Awards or Nobel Prizes are likely to occur. This work serves as a proof of concept that, starting from the level of fundamental knowledge generation, it is possible to detect early signals of applied therapeutic development goals before specialized information from public-facing clinical trials is available.

Here, we show that the information associated with research clusters can provide translation signals even before the later-stage trials are conducted and seminal works are published. The approach used here can flag promising research topics more than 13 years prior to the associated drug approval. The vast majority of research clusters that later contain pivotal studies supporting FDA approvals are identified in advance, most receiving high prediction scores 8 years in advance. In most cases, the prediction signals appear before the associated phase 2 trials appeared.

## Data Availability

Data for Figures 1-7 can be found at Figshare doi: 10.6084/m9.figshare.31937121 (41) in csv format.

## Code availability

Code for Figures 1-7 can be found at Figshare doi: 10.6084/m9.figshare.31937121 (41) as Python Jupyter Notebooks. All packages used (Pandas, Scikit-Learn, Seaborn) are open source and available on PyPI.

## Author Contributions

Conceptualization: BIH; Methodology: SA and BIH; Software: SA and BIH; Validation: SA and BIH; Formal Analysis: SA and BIH; Investigation: SA and BIH; Resources: BIH; Data Curation: SA and BIH; Writing – Original Draft: SA; Writing – Review & Editing: BIH; Visualization: SA and BIH; Supervision: BIH; Project Administration: BIH; Funding Acquisition: BIH.

## Supplementary Figures

**Supplementary Fig 1:**
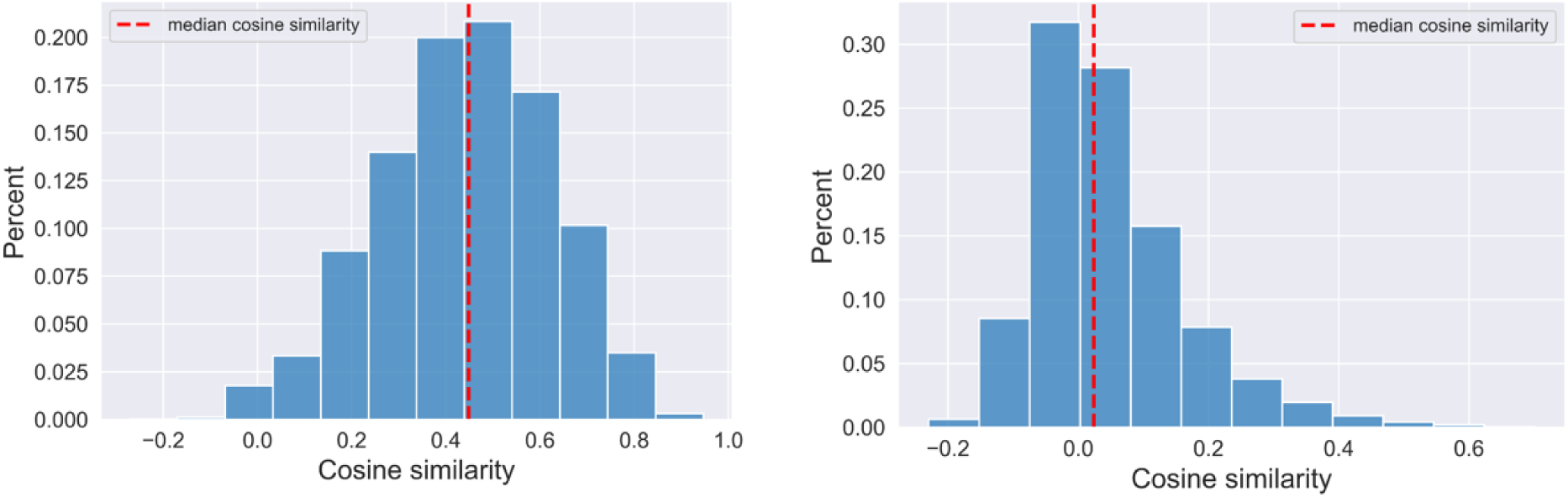
Distribution of cosine similarity of article pairs. (Left) article pairs in the same cluster (right) article pairs from different clusters.

**Supplementary Fig 2:**
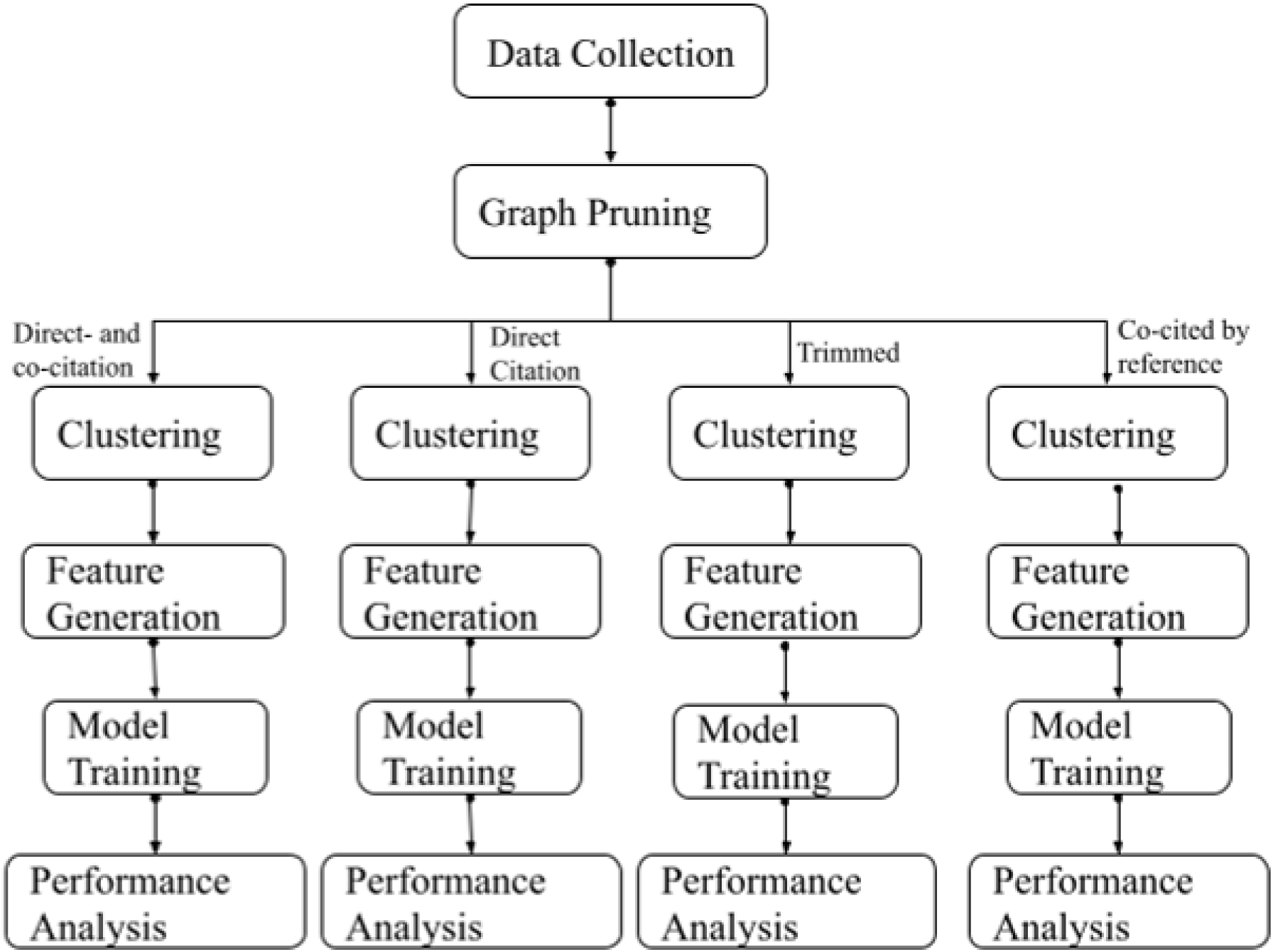
Overview of the classification method using different edge types.

**Supplementary Fig 3:**
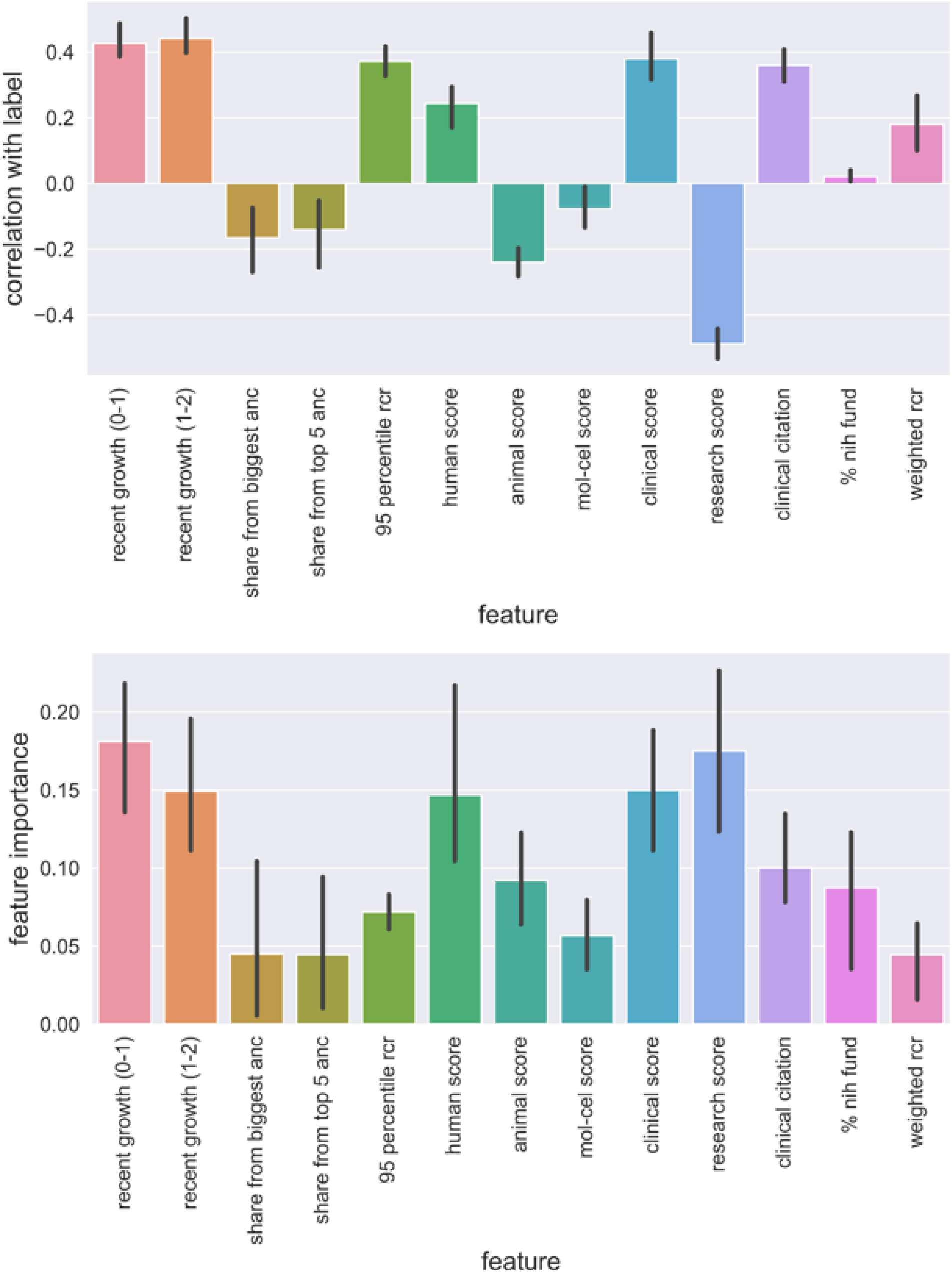
Feature Importance. (Top) top features and their mean Pearson correlation with the label across four edges and 95% CI. (Bottom) mean feature importance of the top features across four edges and 95% CI.

**Supplementary Fig 4:**
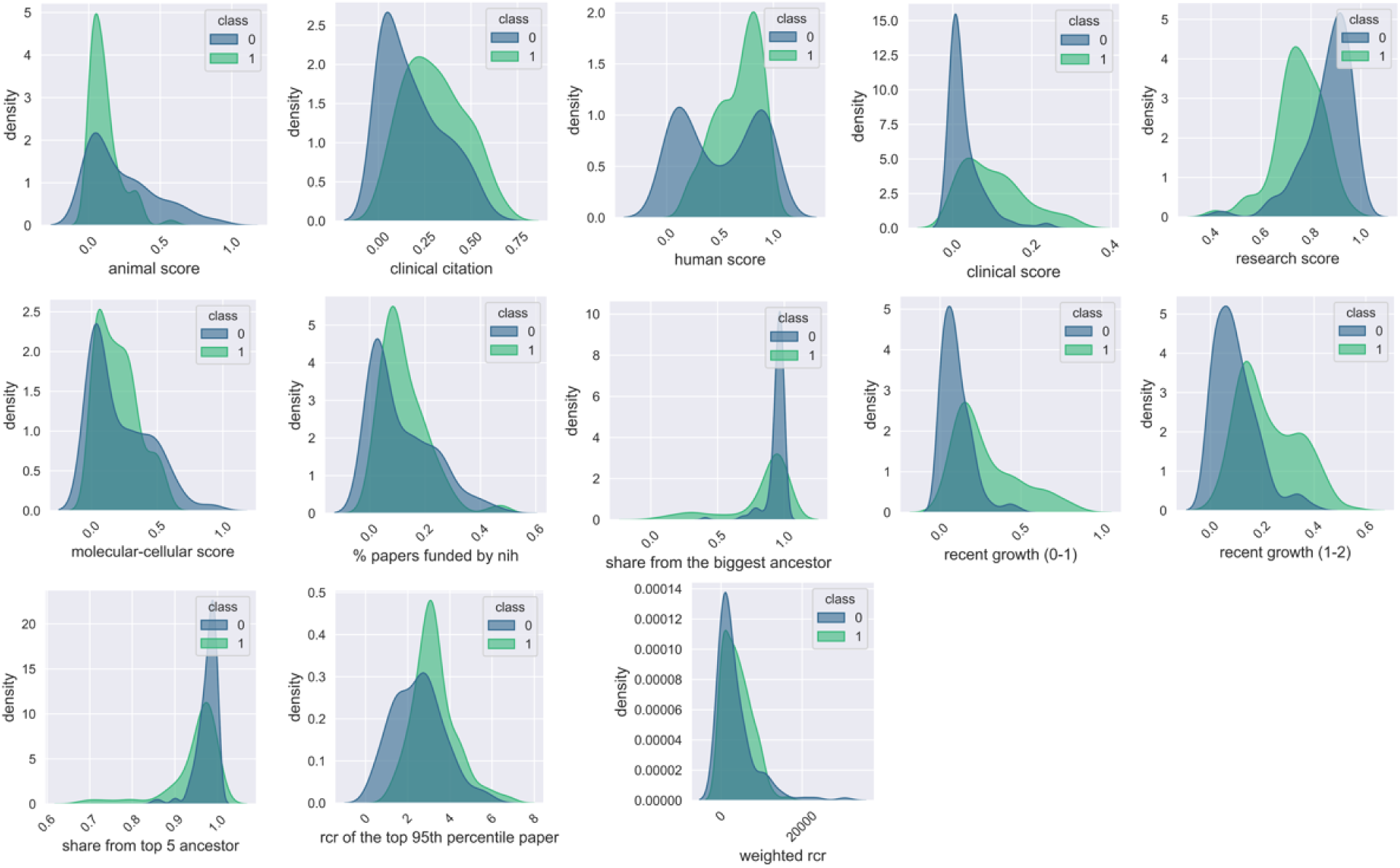
Feature distribution across positive and negative samples. Class 1 (positive) represents RC’s containing pivotal trial publications supporting an FDA-approved drug. Class 0 (negative) represents RC’s with no known link to an FDA-approved drug.

**Supplementary Fig 5:**
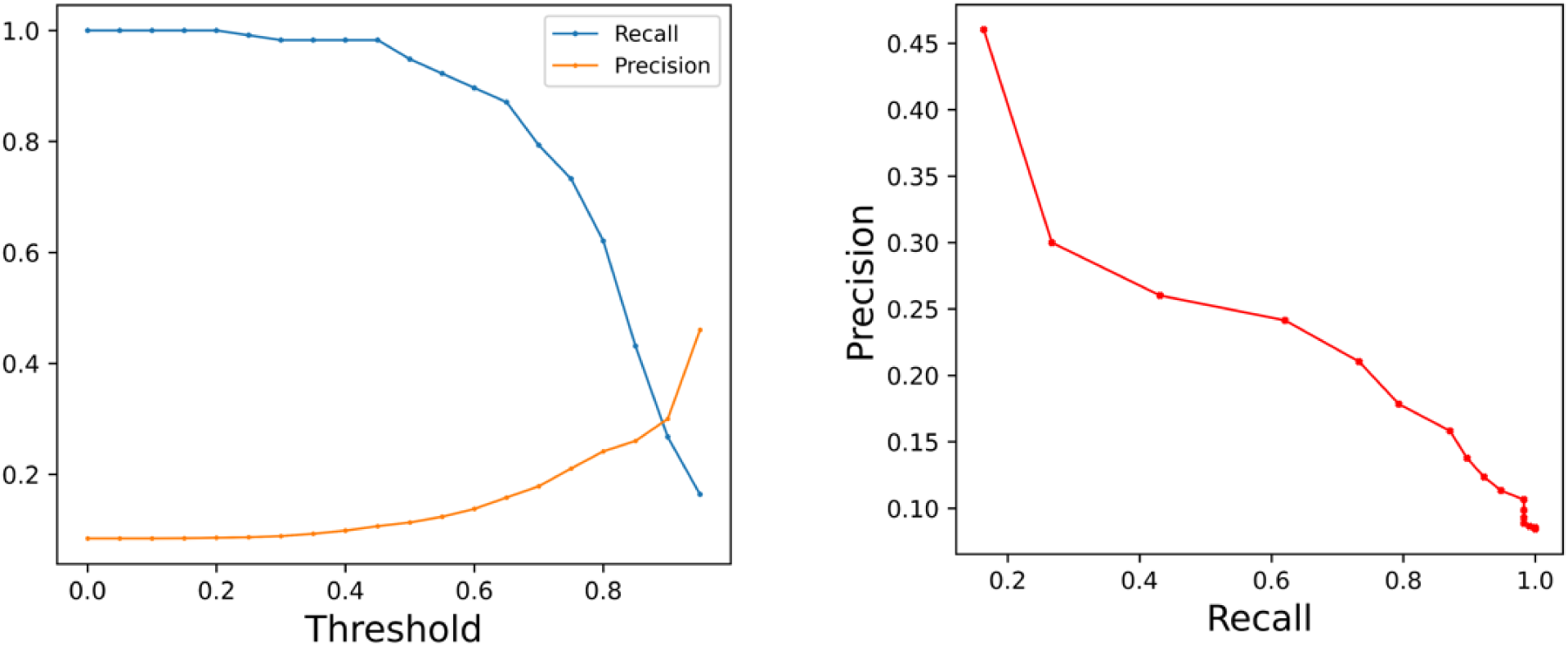
Precision-Recall of the unified model on trimmed edge type. (Left) precision and recall on different thresholds. (Right) precision vs recall graph.

**Supplementary Fig 6:**
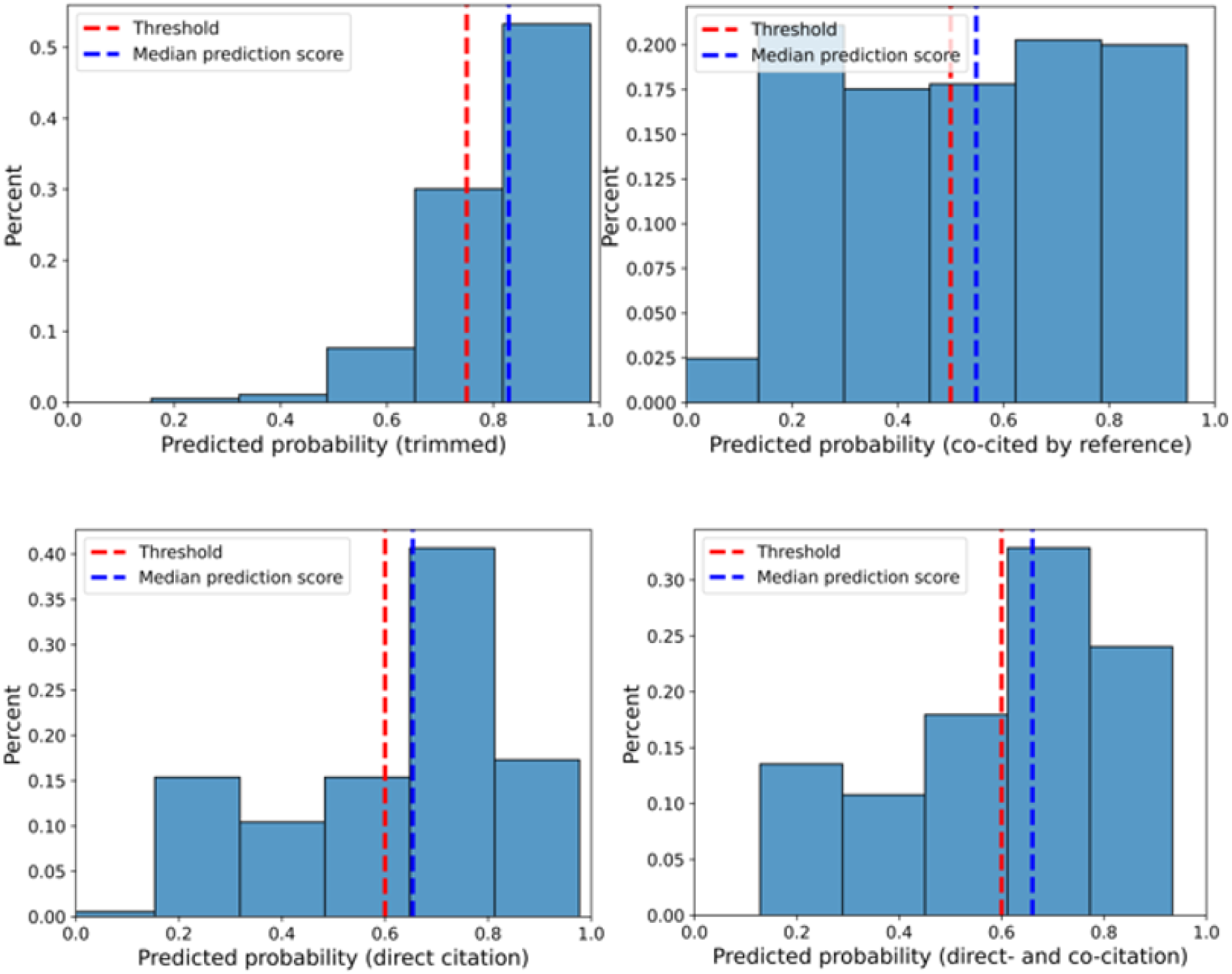
Distribution of prediction score on the exemplar drug clusters for different edge types.

**Supplementary Table 1**: Threshold criteria.

**Supplementary Table 2**: Performance of the unified model on each year from 1978 to 2020.

**Supplementary Table 3**: Performance of the unified model on exemplar drug clusters across different edge types.

**Supplementary Table 4**. 2025 clusters with highest scores for future therapeutics based on the 2023-2025 prediction scores.

