## Supplementary Methods for "Forecasting novel therapeutic development in biomedical research"

### Research Cluster Formation

The graph clustering Leiden algorithm (1) divides the direct citation network into research clusters where articles with similar content and semantic meaning are clustered together. To validate our claim, we randomly selected around 10000 bootstrap article pair samples from the same cluster and around 10000 another article pair samples from different clusters and computed the cosine similarity of the word2vec embedding (2) of their abstracts. The distribution of the cosine similarity is shown in Supplement Fig. 2. We can see that the distribution of cosine similarity of different cluster article pairs is right skewed, with a median similarity around 0.0 whereas the similarity distribution of the same cluster pairs is quite opposite with a median similarity score around 0.4. The figures confirm that the Leiden clustering algorithm is dividing the network into semantically close topics.

### Clustering on different edge types

We prune the direct citation network by removing certain types of edges and generate four different citation graphs. From each citation graph, we create research clusters using the graph clustering Leiden algorithm. For each citation graph and the RCs derived from that, we generate a set of predictive features and build a suitable model using specific features and hyperparameters tuned for the specific edge type. The overview of our approach is shown in Supplemental Fig. 2. To evaluate the performance or our approach, we collect data about a set of drug therapeutics and the pivotal clinical trial associated with the drugs and map the publication associated with the pivotal trials to the specific clusters containing those publications to generate training and test sets for each edge type. We test the performance of the four models on the four edge types (Table 1). Finally, we built a unified model using the same set of features and test the performance of all four different network structures and the clusters generated from them (Supplement Table 2). As the trimmed edge type outperforms the rest of the edges, we conduct our further time series validation on exemplar and blockbuster drugs on the trimmed edge type.

### Feature Selection

We use scikit-learn package (mutual_info_classif) for mutual information estimation for classification to explore the importance of various features (Supplemental Figs 3 and 4). To resolve multicollinearity and reduce model complexity, we exclude features that are weakly correlated to the label and also the ones that are strongly correlated to the other features. The feature set falls into five broad categories based on the type of information they provide.

The first set of features is intended to capture the emerging publication patterns in a research cluster. It measures the recent explosion in the number of research publications in a particular cluster. These features capture the recent scholarly attention achieved by the research topic associated with the cluster from the research community and a previous study reported the contribution of similar features in predicting intervention approval (3). Higher recent share in the number of publications in a particular cluster indicates the growing interest of scientists and researchers in the small sub-topic of science associated with the cluster. It provides a social signal of an exciting and important prospect in that line of research. Research funding organizations are usually interested in supporting such research. We have measured the growth of publication in a cluster in the terminal year and the year before that and both features are positively correlated with the label.

The second set of features measures the focus of a research cluster toward the human, animal, and molecular-cellular axis in the triangle of biomedicine (4). The triangle of biomedicine places publications across human, animal, and molecular/cellular axes based on their Medical Subject Headings (MeSH). These features impact the translational process and hence, are being used in predicting bench-to-bedside translation (5). The position of a research topic evolves over the years and topics with translational attributes are expected to gradually move from the animal or molecular/cellular corner towards the human corner. Clinical trials are generally closer to the human vertex, and they become gradually closer with the increasing phase of the trials (4). In Supplemental Fig 3(a), we can see that the label has a significant positive correlation with the degree of human and a negative correlation with the degree of animal and molecular cellular biology.

The third set of features measures the clinical orientation of the articles within the clusters. Clinical citation count provides strong evidence of clinical impact in biomedical literature (5). Having a higher number of clinical articles as well as a higher proportion of clinical citations indicate the potential of a field of science in generating clinical research. These features are positively associated with a greater likelihood of drug development. On the other hand, having a higher proportion of research articles may indicate that the cluster is basic research-oriented or still in an early stage.

The fourth set of features measures the scholarly influence and funding status of the articles within a particular cluster. We exploit the information about the proportion of highly and moderately influential articles in a particular cluster. The influence is measured using the National Institute of Health's (NIH) field and time-normalized, article-level metric Relative Citation Ratio (6). RCR measures the influence of a research article while reducing the impact of field and time on citation count. We also consider the proportion of NIH-funded articles in the clusters as previous studies showed that research funding impacts the research outcome and increases research productivity (7).

The last set of features measures how a particular line of research evolved over the past few years. We measure fragmentation and discontinuity in the trajectory of the cluster. Interestingly, the feature correlation suggests that fragmentation and discontinuity of an existing topic strongly signal a potentially successful drug outcome. We have two features in this context. While one feature indicates the continuation of an existing research topic, and another one indicates less fragmentation of an existing research topic in recent years.

### Building the unified model

We develop a unified model to compare the performances of different edge types in similar settings. We tested the performance of the unified model on the clusters for each year from 1978 to 2020 for the four edge types. To test our model on a particular year's entire set of clusters, we held out all instances of that particular year from the training data during the training and made predictions on the clusters of that year. While testing the model on the entire set of clusters for each year, we applied a set of cut-off criteria to discard unstable or less promising clusters for drug intervention. The criteria were selected in a way to discard the clusters that are far below compared to the positive drug clusters in the training data in terms of the 10-percentile value of every important feature from our consideration. The accuracy of prediction is calculated as the proportion of true positive samples detected as positive by the model. We considered models' prediction probabilities greater than 0.75, 0.60, 0.50, and 0.60 as positive predictions for trimmed, direct citation, co-cited by reference, and direct- and co-citation edge types sequentially. The thresholds were selected based on the prediction score distributions across 4 edge types to include at least half of the exemplar clusters in the positive category for all edges.

### Threshold Criteria

A threshold criterion is adopted to remove unpromising and unstable clusters from consideration. Unpromising in this context means clusters that are less likely to generate drug therapeutics. In order to find such clusters, we turned to the features that looked distinctive across the positive and negative clusters. We removed the clusters having feature values even lower than the mean 10th percentile value of positive training clusters for the four edge types. The criteria we used to remove clusters from our consideration are listed in Supplemental Table 1.

We found these criteria to be effective in defining research clusters with an adequate combination of basic and applied focused research as well as in defining stable clusters with influential research. Also, it helps to remove unstable clusters and reduce false positives while making predictions on all the clusters of a particular year.

### Model Testing

To test our model on a particular year's clusters, we held out all instances of that particular year from the training data during the training and made predictions on the clusters of that year. This approach not only lets us test our model's capacity to predict true positive clusters but also provides insight into the yearly false positive prediction rate given by the model. Also, concerns may arise about the predictions on year "X" while the model is using data from year "X+1", "X+2" or so regarding the possible impact of lookahead bias. However, the model does not use any identifiable information such as cluster id, year, etc as features that can create data leakage in the model.

Supplemental Table 2 shows the accuracy, false positive rate, and other performance metrics of the unified model on four edge types. Trimmed edge leads the accuracy and cross-validated f1 score over the other three. While direct and co-cited by reference edges provide fewer false positives, their accuracy is much lower than trimmed edge type. Moreover, this table provides valuable insight into the interaction between network pruning and the machine learning model. Although we started with the same citation network, the pruning strategy creates a big difference in various performance measures. It is indeed interesting to see that the local network structure makes significant impacts on how much information is accessible to the machine learning model for making predictions. Even though the network with trimmed edge type has the least number of edges which makes it much smaller compared to the largest network representation (direct), it significantly overperformed the direct edge and also the other two.

The precision-recall graph of the unified model on the trimmed edge type is shown in Supplement Fig 5. The lack of information regarding hundreds of drugs in the dataset increases the number of false positives which is further reflected in the lower precision curve. Due to the having data for only a subset of drugs, it is possible that some of the false positives are indeed true positives that are not included in our dataset. However, it was not feasible to investigate each false positive instance manually. Therefore, the false positive rate should be considered with caution because they can still produce drug breakthroughs or can be out-of-sample true positives.

### Testing Unified model on the exemplar drug clusters

We tested the performance of the unified model on 370 exemplar drug clusters for different edge types (Supplement Table 3). The prediction score distribution of the positive clusters largely varies across different edge types (Supplement Fig. 6).

We analyzed the performance of the unified model across the 15 years trajectory of the 99 exemplar drugs to analyze how early our model can predict the emergence of a drug relative to its approval year. We record the prediction score given by the model across the whole trajectory of each drug and identify the first occurrence of a higher prediction (>0.75) score in the trajectory.

The result of the analysis is shown in Fig. 3. It was interesting to see that approximately 75% of the exemplar drugs were identified as promising by our model 9 to 1 year before the approval of the associated drugs. In fact, 54.5% of the drugs were identified during the timeline of 4-6 years before approval. While four drugs were identified after their approval, our model failed to identify the emergence of 14 drugs as no higher prediction score appeared throughout their trajectories.
